# DeEPsnap: human essential gene prediction by integrating multi-omics data

**DOI:** 10.1101/2024.06.20.599958

**Authors:** Xue Zhang, Weijia Xiao, Brent Cochran, Wangxin Xiao

## Abstract

Essential genes are necessary for the survival or reproduction of a living organism. The prediction and analysis of gene essentiality can advance our understanding of basic life and human diseases, and further boost the development of new drugs. Wet lab methods for identifying cell essential genes are often costly, time-consuming, and laborious. As a complement, computational methods have been proposed to predict essential genes by integrating multiple biological data sources. Most of these methods are evaluated on model organisms. However, prediction methods for human essential genes are still limited and the relationship between human gene essentiality and different biological information still needs to be explored. In addition, exploring suitable deep learning techniques to overcome the limitations of traditional machine learning methods and improve prediction accuracy is also important and interesting. We propose a snapshot ensemble deep neural network method, DeEPsnap, to predict human essential genes. DeEPsnap integrates sequence features derived from DNA and protein sequence data with features extracted or learned from multiple types of functional data, such as gene ontology, protein complex, protein domain, and protein-protein interaction network. More than 200 features from these biological data are extracted/learned which are integrated together to train a series of cost-sensitive deep neural networks by utilizing multiple deep learning techniques. The proposed snapshot mechanism enables us to train multiple models without increasing extra training effort and cost. The experimental results of 10-fold cross-validation show that DeEPsnap can accurately predict human gene essentiality with an average AUROC (Area Under the Receiver Operating Characteristic curve) of 96.1%, the average AUPRC (Area under the Precision-Recall curve) of 93.82%, the average accuracy of 92.21%, and the average F1 measure about 80.62%. In addition, the comparison of experimental results shows that DeEPsnap outperforms several popular traditional machine learning models and deep learning models. We have demonstrated that the proposed method, DeEPsnap, is effective for predicting human essential genes.

## Introduction

Human beings have more than 20,000 genes, which form a redundant and highly fault-tolerant system. Among these genes, some are vital for the survival and reproduction of us, but others are not. These two groups of genes are named as essential genes and nonessential genes. Essential genes are a group of fundamental genes necessary for a specific organism to survive in a specific environment. Cell essential genes refer to a subset of genes that are indispensable for the viability of individual human cell types [1, 2] as opposed to genes required for the survival of a multicellular organism. Here we focus on predicting cell essential genes. These cell essential genes encode conservative functional elements which mainly contribute to DNA replication, gene translation, gene transcription, and substance transportation. The identification and analysis of essential genes is very important for understanding the minimal requirements of basic life, and it’s vital for drug-target identification, synthetic biology, and cancer research.

There are two ways to identify essential genes, wet lab experimental methods and computational methods. For example, gene direct deletion and transposon-based randomized mutagenesis have been used to identify essential genes for bacteria and yeast in the genome scale [3]; microinject KO and nuclear transfer techniques have been used to identify essential genes in mice [4]; the CRISPR/Cas9 genome editing system has been used to identify essential genes from human cell lines [1,2,5]. Experimental methods are often costly, time-consuming, and laborious. The accumulation of essential gene datasets and the sequence data as well as multiple functional data enables researchers to explore the relationships between gene essentiality and different genomics data and to develop effective models to predict gene essentiality. These computational methods can greatly reduce the cost and time involved in finding essential genes which further boosts our understanding of basic life and human diseases, and helps quickly find new drug targets and develop drugs.

Computational methods can be classified into two groups. One focus is to design a centrality measure to rank proteins/genes, while the other focus is to integrate multiple features using machine learning to predict gene essentiality. The most widely used centrality measures include degree centrality, betweenness centrality, closeness centrality, and eigenvector centrality, to name a few. These centrality measures have been found to have a relationship with gene essentiality in multiple model organisms and humans [6, 7], however, they can only differentiate a subset of essential genes from nonessential genes. One reason might be the incompleteness and false positive/false negative interactions in the protein-protein interaction (PPI) networks, the other reason might be the fact that gene essentiality relates to multiple biological factors rather than only the topological characteristics. Due to these reasons, researchers have proposed some more complicated centrality measures by integrating network topological properties with other biological information. For example, Zhang et al. proposed a method, CoEWC, to capture the common properties of essential genes in both date hubs and party hubs by integrating network topological properties with gene expression profiles, which showed significant improvement in prediction ability compared to those only based on the PPI networks [6]. An ensemble framework was also proposed based on gene expression data and the PPI network, which can greatly enhance the prediction power of commonly used centrality measures [8]. Luo et al. proposed a method, LIDC, to predict essential proteins by combining local interaction density with protein complexes [9]. Zhang et al. proposed OGN by integrating network topological properties, gene expression profiles, and orthologous information [10]. GOS was proposed by Li et al. by integrating gene expression, orthologous information, subcellular localization, and PPI networks [11]. These proposed integrated centrality measures have improved prediction power over the ones only based on PPI network topological properties. However, they still have limited prediction accuracy since gene essentiality relates to many complicated factors that are impossible to represent by a scalar score. At this point, machine learning is a good choice to fully utilize multiple features for predicting essential genes.

Many machine learning models and deep learning frameworks have been proposed and successfully applied in different fields. For example, density-based neural network was used for pavement distress image classification [12]; convolutional neural network (CNN) was used for drug-target prediction [13]; active learning and transductive k-nearest neighbors were used for text categorization [14, 15]; Support Vector Machines (SVM) was used for essential gene prediction [16]; graph attention networks (GAT) was used to predict essential genes [17], drug-induced liver injury [18] and cardiotoxicity related hERG channel blockers [19]; Auto-encoder was used to predict metabolite-disease associations [20]. In the research field of essential gene prediction, many machine learning-based prediction methods have been proposed to integrate features from multiple omics data [21]. As shown in the review article [21], traditional machine learning methods were used to predict gene essentiality, and most of them were evaluated on data from model organisms. In these methods, topological features together with features from sequence and other functional genomics data were extracted manually and then used to train the models. How to extract informative features is very important and challenging, which requires ample domain knowledge as well as the understanding of what relationship exists between gene essentiality and each omics data. Usually, we only know that an omics data would contribute to gene essentiality, but we don’t know what attributes of it have such an effect and how to represent it. This puts a limitation on traditional machine learning methods to obtain good prediction accuracy.

In recent years, deep learning techniques have been used to automatically extract features and to train a more powerful classification model for predicting essential genes. Grover et al. proposed a network embedding method based on deep learning, node2vec, to learn a low-dimension representation for each node [22]. This method has been used to extract topological features from PPI networks for predicting essential genes, and these features are more informative than those obtained via some popular centrality measures [7, 23, 24, 25]. CNN was used to extract local patterns from time-serial gene expression profiles from S. cerevisiae [23] and Zeng et al. also used bidirectional long short-term memory (LSTM) cells to extract features from the same gene expression profiles [24]. The automatically learned features are combined with other manually extracted ones to train a deep learning model for human essential gene prediction [7]. A six-hidden-layer neural network was designed to predict essential genes in microbes by only using manually extracted features from sequence data [26]. Li et al. proposed a deep learning method for predicting cell line-specific essential genes based on sequence data [27], which integrates a convolutional neural network, bidirectional long short-term memory, and a multi-head self-attention mechanism together expecting to learn short- and long-range information from protein sequence and to provide residue-level model interpretability. Since sequence data is not cell line-specific, the model can only be trained and tested for each cell line separately. Yue et al. proposed a deep learning model for predicting essential proteins by integrating the PPI network, subcellular localization, and gene expression profiles together, which is evaluated on the data of Saccharomyces cerevisiae [28].

Recently, human essential genes have been identified in several human cancer cell lines by utilizing CRISPR-Cas9 and gene-trap technology [1,2,29]. These identified essential genes provide a clear definition of the requirements for sustaining the basic cell activities of individual human tumor cell types, and can be regarded as targets for cancer treatment. These essential gene datasets together with other available biological data sources enable us to test one important and interesting assumption that human gene essentiality might be accurately predicted using computational methods. In this paper, we propose a Deep learning-based Essential Protein prediction method using a novel snapshot ensemble mechanism, DeEPsnap, to predict human essential genes. DeEPsnap integrates features from five omics data, including features derived from nucleotide sequence and protein sequence data, features learned from the PPI network, features encoded using gene ontology (GO) enrichment scores, features from protein complexes, and features from protein domain data. The proposed snapshot ensemble mechanism is inspired by the work of [30], which shows that cyclic learning rates can be effective for training convolutional neural networks. In this paper, we propose a new cyclic learning method for our essential gene prediction problem. We show that DeEPsnap can accurately predict human essential genes. The main contributions of this paper include: 1) extract useful features from multi-omics data and integrate them for predicting human gene essentiality. 2) propose a new snapshot ensemble mechanism to improve the prediction performance. 3) showcase the usefulness and effectiveness of features extracted from five omics data to gene essentiality prediction and the different contributions of the extracted features from different omics data.

## Snapshot ensemble deep learning model

DeEPsnap consists of three main modules: input data module, feature extraction and feature learning module, as well as classification module. The flowchart of DeEPsnap is shown in Fig 1.

**Fig 1.**
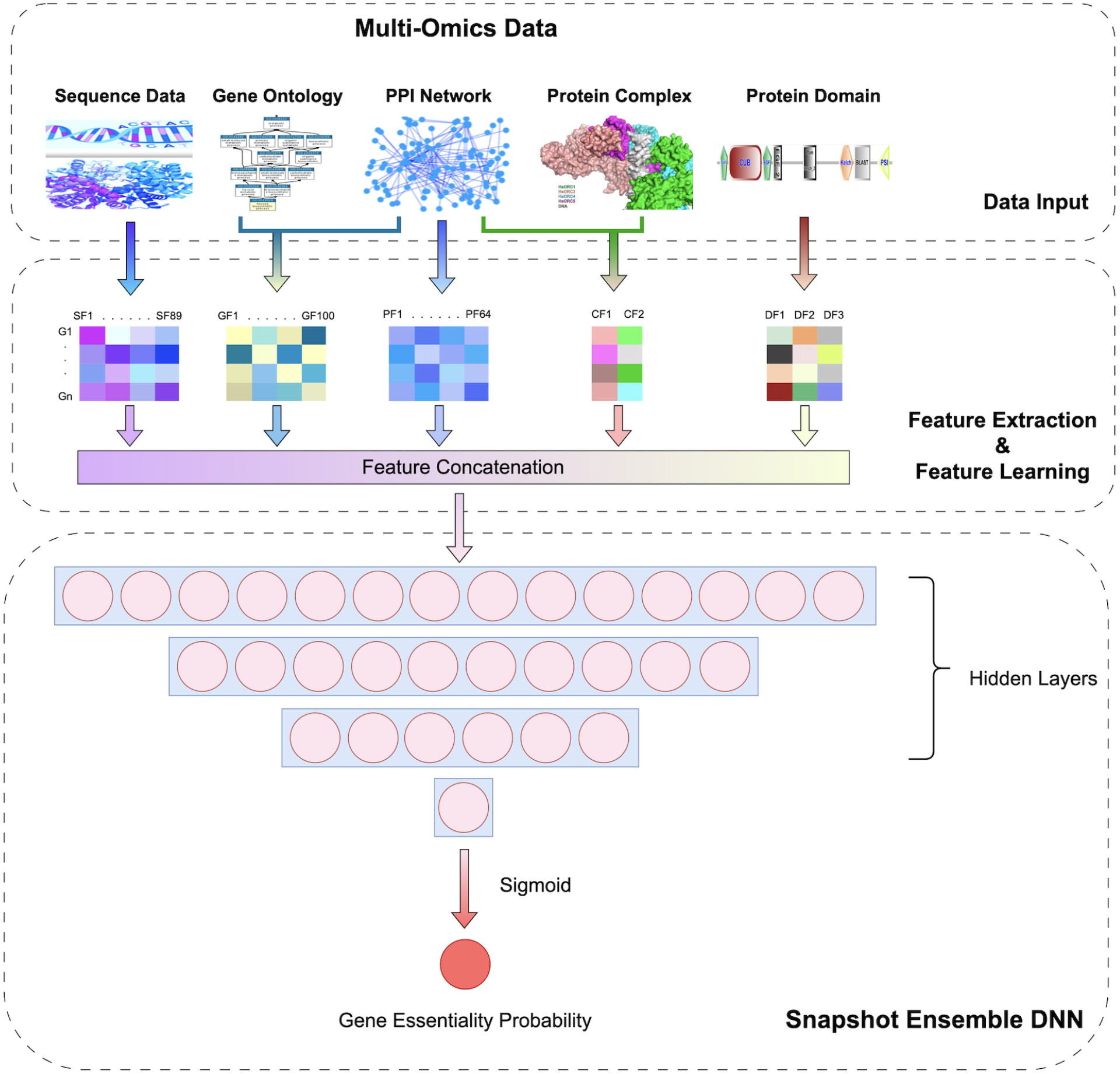
The Flowchart of DeEPsnap.

### Input data module

In this paper, we mainly consider five omics data to explore their relevance and efficiency for predicting gene essentiality. As shown in Fig 1, the input data module includes five biological data sources, which are sequence data (DNA sequence and protein sequence), gene ontology, protein-protein interaction network (PPI network), protein complex, and protein domain.

### Feature extraction and feature learning module

The feature extraction and feature learning module uses different methods to extract or learn features from multiple omics data. Then the features are concatenated together as input to the classification module. In this paper, we mainly consider five types of features. More features can be easily integrated into our model to potentially further improve prediction accuracy.

#### GO enrichment score encoded features

Features encoded with gene ontology (GO) enrichment scores are calculated as follows. We choose the first 100 GO terms from the cellular component (CC) subcategory to encode the genes, where CC terms are ranked in descending order based on the number of essential genes involved in the terms. For each gene, we first obtain its direct neighbors from the PPI network to form a gene set consisting of this gene and its neighbors, then do gene ontology enrichment analysis for this gene set against the 100 GO terms using a hypergeometric test. The enrichment score is calculated as –log10(p-value) for each GO term. In this way, we get a 100-dimension feature vector for each gene. The GO enrichment score encoded features capture information from both the PPI network and subcellular localization of genes.

#### Features derived from protein complex data

From protein complexes, we extract two features for each gene. The first feature is the number of protein complexes a gene involved in. The second feature is calculated as follows. For a gene, we first get the gene set N consisting of its direct neighbors in a PPI network. Suppose there are M neighbors of this gene involved in a protein complex. We calculate a score s = |M|/|N| as the ratio of its neighbors involved in a protein complex. The second feature is the sum of the ratios across all the protein complexes. Therefore, the second feature also considers the information from the PPI network in addition to that from the protein complex data.

#### Features learned from the PPI network

Network features are learned based on a network embedding method, node2vec [22]. Each gene is represented by a 64-dimension feature vector learned from the PPI network. Previous studies showed that this low-dimension representation learned using node2vec is superior to the features calculated by popular centrality measures [7, 23, 24].

#### Features derived from sequence data

Sequence features consist of codon frequency, maximum relative synonymous codon usage (RSCUmax), codon adaptation index (CAI), gene length, GC content, amino acid frequency, and protein sequence length. There are 89 sequence features in total. For more details about how these features are calculated, please refer to [7].

#### Features derived from protein domain data

From protein domain data, we extract three features for each gene. The first feature is the number of domain types a protein has. The second feature is the number of unique domain types a protein has. The third feature is the sum of inverse domain frequency (IDF). The frequency *f* of a domain is the number of proteins that have this domain. Its IDF is 1/*f*. Suppose a protein *u* has *n* domains, then the third feature of *u* is calculated as in (1).

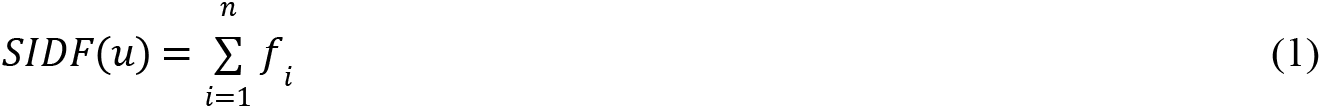

### Classification module

#### Baseline model

The classification module of DeEPsnap is based on a snapshot ensemble deep neural network. The baseline model here is a multilayer perceptron enhanced by several deep-learning techniques, and we name it DNN. It includes one input layer, three hidden layers, and one output layer. We use the rectified linear unit (ReLU) as the activation function for all the hidden layers, while the output layer uses the sigmoid activation function to perform discrete classification. The loss function in DeEPsnap is binary cross-entropy, as shown in (2), where *N* is the number of samples. Each hidden layer is followed by a dropout layer to make the network less sensitive to noise in the training data and to increase its generalization ability. To address the imbalanced learning issue inherent in the essential gene prediction problem, we utilize class weight to train a weighted neural network, which gives larger penalties when the model misclassifies an instance from the minority class. The use of class weight encourages the model to pay more attention to instances from the minority class than those from the majority class leading to a more balanced and effective classifier.

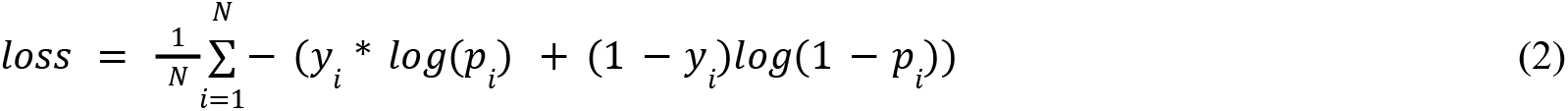

#### Cyclic sine annealing

In machine learning, ensemble methods utilize multiple learning algorithms to obtain better predictive performance than could be obtained from any of the constituent learning algorithms alone. Likewise, ensembles of deep neural networks are known to be much more robust and accurate than individual networks. However, training multiple deep neural networks is computationally expensive. In this paper, we propose a snapshot ensemble mechanism to obtain multiple trained models without incurring extra training costs. The snapshot ensemble mechanism creates an ensemble of accurate and diverse models from a single training process. We expect such an optimization process that visits several local minima before converging to a final solution. Model snapshots at these various minima are taken and their predictions are averaged at the test time.

To converge to multiple local minima, we propose a cyclic annealing scheduler based on the sine function. The learning rate is decreasing at a very fast pace, encouraging the model to converge towards its first local minimum after as few as 5 epochs in the DeEPsnap training process. The optimization is then continued at a larger learning rate in order to perturb the model and dislodge it from the minimum. This process is repeated several times to obtain multiple convergences. Formally, the learning rate *lr*(*t*) has the form shown in (3), where *t* is the current training epoch number (0-based), *C* is the total number of epochs in an annealing cycle, *lr*_0_ is the initial learning rate at the beginning of each cycle, while % represents the modulo operation. If the number of the total training epochs is set as *T*, then the training process is splitted into *M* = ⌊*T*/*C*⌋ cycles. If *T* is not divisible by *C*, the remaining epochs will still be trained but will not contribute to the final ensemble model. Each cycle starts with a larger learning rate, which is annealed gradually to a smaller learning rate. Finally, the *M* models will be saved and used to make predictions for the testing stage. Fig 2 illustrates the learning rate cycles in DeEPsnap and the snapshot models. In each learning rate cycle, the learning rate starts from 0.001 and anneals towards 0. The intermediate models, denoted by the dotted lines, form an ensemble at the end of training. The parameters of DeEPsnap in our experiments are shown in Table 1.

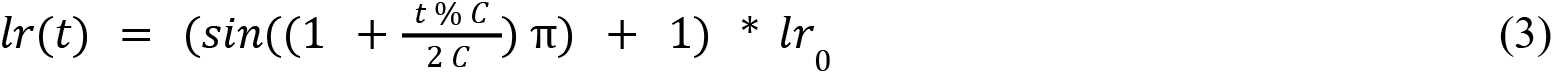

**Fig 2.**
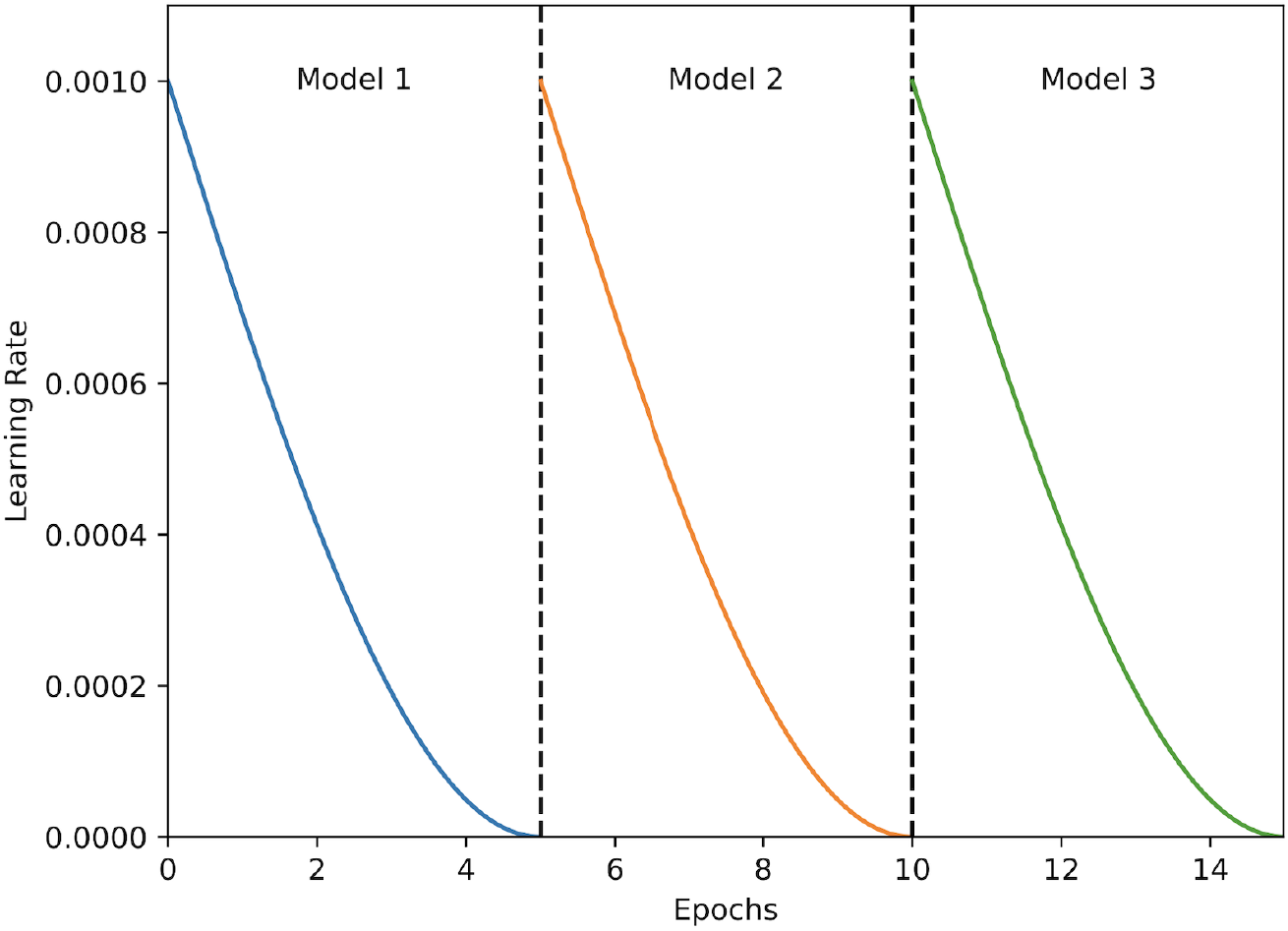
Learning rate cycles and snapshot ensemble models.

**Table 1.**
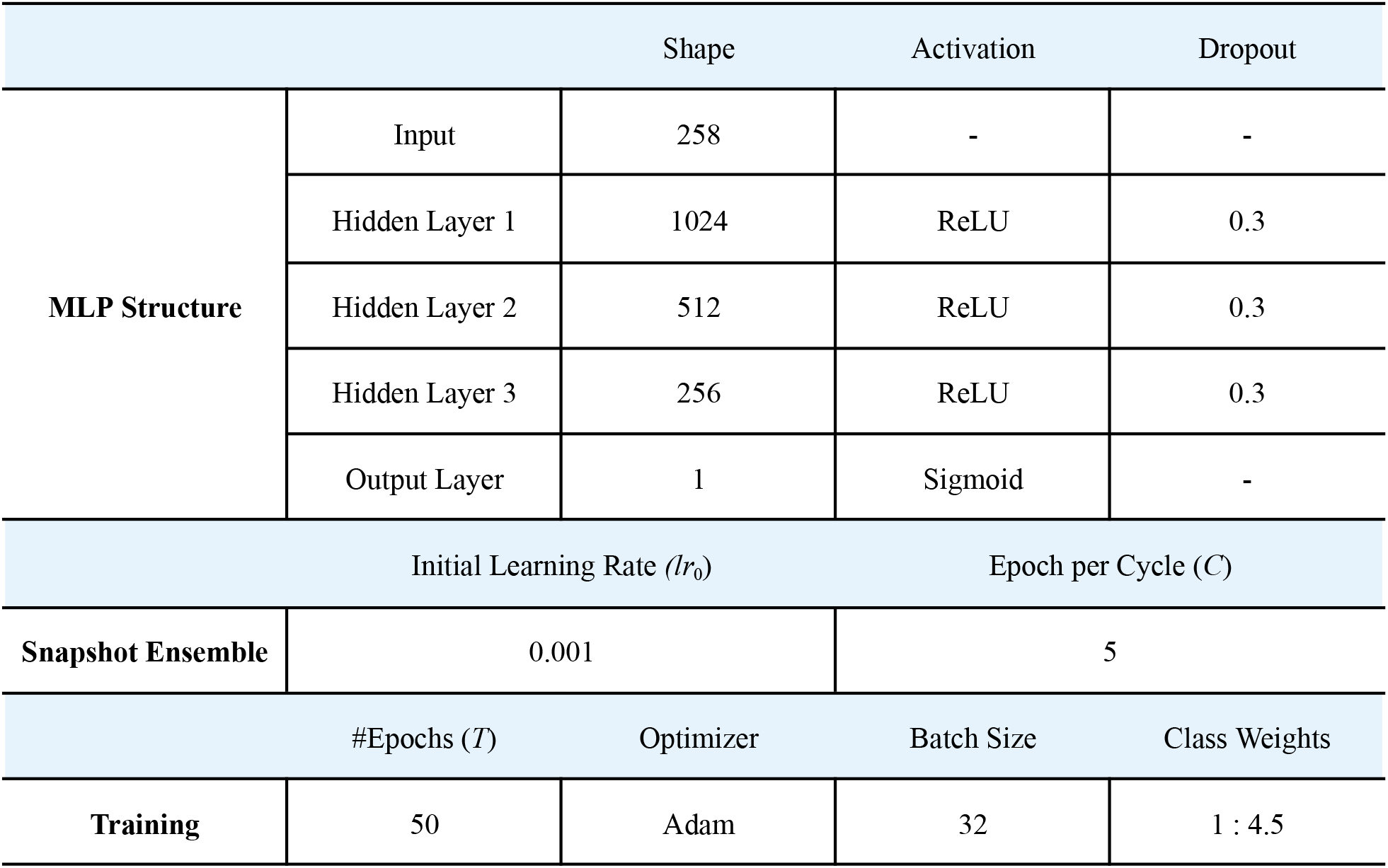
Parameters of DeEPsnap.

## Results and discussion

### Datasets

Human essential genes are downloaded from the DEG database [31]. There are 16 human essential gene datasets among which 13 datasets are from [1,2,29]. We chose the genes contained at least in 5 datasets as our essential gene dataset. Excluding the genes annotated as essential genes in DEG, the other genes are considered nonessential genes.

The DNA sequence data and protein sequence data are downloaded from Ensembl [32] (release 97, July 2019). We download the PPI data from BioGRID [33] (release 3.5.181, February 2020). Only physical interactions between human genes are used. After filtering out self-interactions and several small subgraphs, we obtain a protein-protein interaction network with 17,762 nodes and 355,647 edges. This interaction network is used to learn embedding features for each gene. It’s also used in aiding to compute some features from gene ontology, protein complex, and protein domain.

Gene Ontology data are downloaded from the Gene Ontology website [34,35] and protein complex data are downloaded from CORUM [36]. Protein domain data are from the Pfam database [37], and we collect this data via the Ensembl BioMart [32]. The genes having sequence features, network embedding features as well as GO enrichment scores are used for the following classification performance evaluation. In total 2009 essential genes and 8414 nonessential genes are used for the following analysis.

### Evaluation metrics

We use multiple metrics to evaluate the performance of DeEPsnap. The first metric is the area under the receiver operating characteristic (ROC) curve (AUROC). ROC plot represents the trade-off between sensitivity and specificity for all possible thresholds. The second metric is the area under the precision-recall curve (AUPRC). Precision-recall (PR) curves summarize the trade-off between the true positive rate and the positive predictive value using different probability thresholds. ROC curves are appropriate for balanced classification problems while PR curves are more appropriate for imbalanced datasets. Since essential gene prediction here is an imbalanced classification problem, the AUPRC metric is more important than AUROC. The third metric is the balanced accuracy (BA), and the fourth metric is the F1 measure. In addition to these four comprehensive metrics, we also give the comparison results in terms of accuracy. The definitions of BA, accuracy, and F1 are defined in (4) - (6), where *TP*, *TN*, *FP*, and *FN* are the number of true positives, true negatives, false positives, and false negatives, respectively.

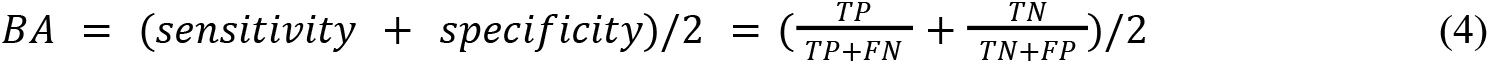

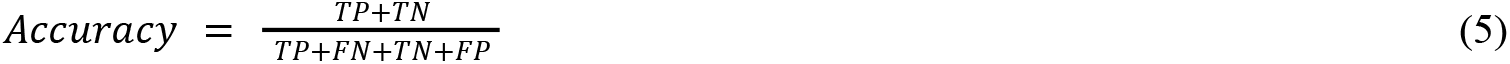

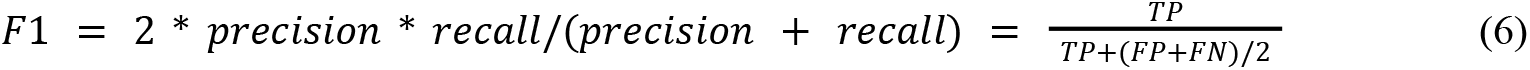

### Performance evaluation

In the following experiments, we use the parameters for DeEPsnap as shown in Table 1. In order to cope with the imbalanced learning issue, we set the class weight to 4.5 for essential genes and 1 for nonessential genes. The stratified randomized 10-fold cross-validation is used to evaluate the performance of DeEPsnap. At each fold, 10% data are held out for testing, and 90% data are used for training.

Fig 3 presents the ROC curve of the DeEPsnap across the 10-fold cross-validation. From Fig 2 we can see that DeEPsnap reaches its best performance at folds 5,7 and 9 with AUROC = 0.97. The average AUROC of DeEPsnap across the 10-fold cross-validation is 96.1% with standard deviation STD = 0.65%, and the average AUPRC is 93.82% with STD = 1.13%. In addition, the performance of DeEPsnap is quite stable since the difference is less than 2.12% between its best and worst AUROC scores across the 10-fold cross-validation. The worst AUPRC score is still above 91.8% which indicates that DeEPsnap is very effective for predicting human essential genes. In addition to good scores of AUROC and AUPRC, its average accuracy, BA, and F1 scores are 92.21%, 89.09%, and 80.62% respectively.

**Fig 3.**
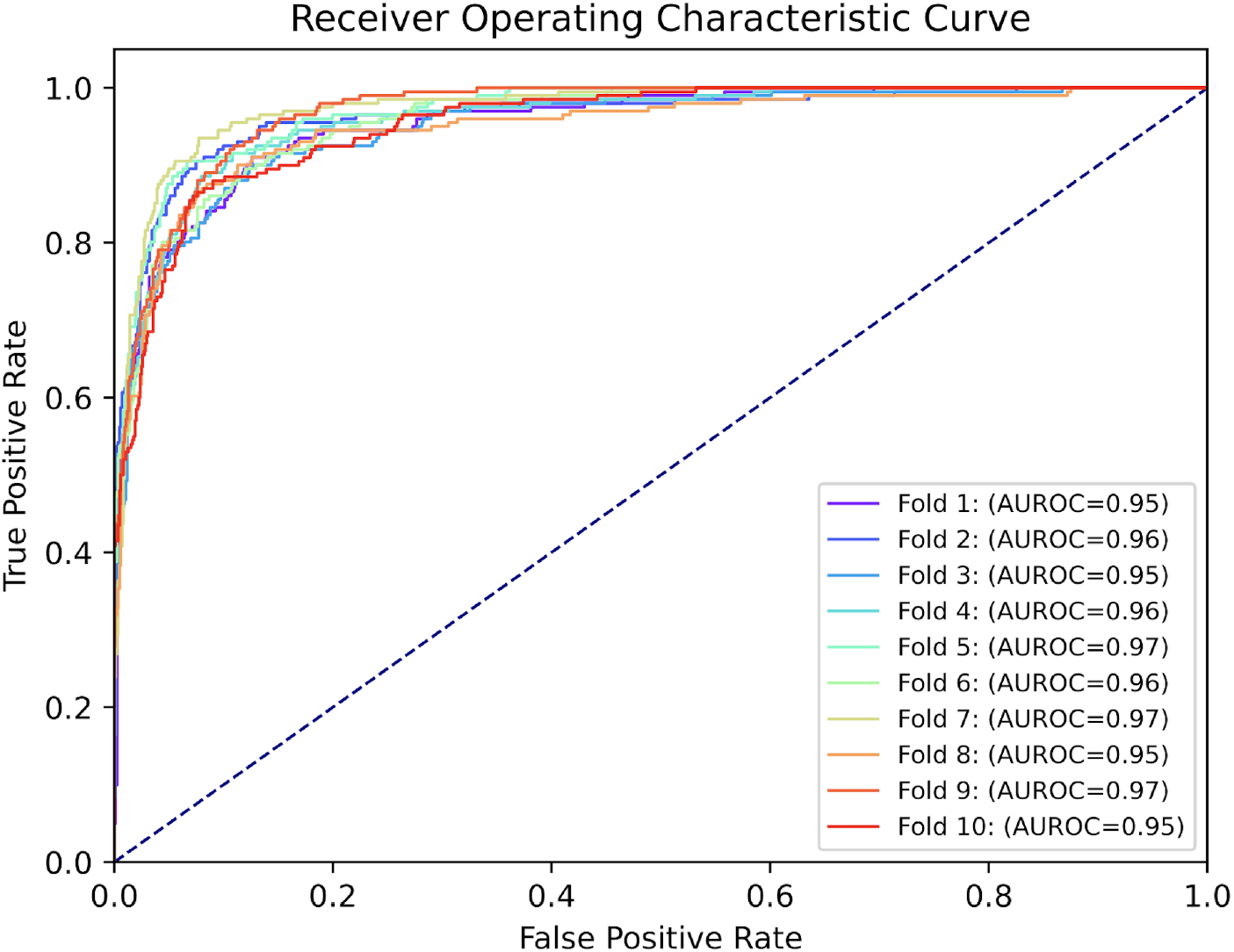
The ROC curve of DeEPsnap.

### Performance comparison with other machine learning models

In order to demonstrate the superiority of DeEPsnap, we also compare it with three popular traditional machine learning models—SVM, Random Forest (RF), and Adaboost—as well as a recent deep learning-based essential gene prediction model DeepHE, and two deep learning models (GAT and DNN). For a fair comparison, DeEPsnap, the three traditional machine learning models, and DNN use the same input features so the only difference here is the classification method. GAT is based on the implementation and parameter settings in [17], while we use an early-stopping mechanism (patience = 200 epochs, 10% training data are used as validation data) to save the best model for the testing stage and set the maximum epochs = 3000. Since GAT is a graph neural network that can capture the structure information of input graphs, we use the PPI network as the input graph and the features extracted from the other four omics data as node features. DNN is the baseline model used in DeEPsnap, while it uses a constant learning rate of 0.001 and an early stopping mechanism with patience of 15 epochs (10% training data are used as validation data). For DeepHE, we use the features and model structures presented in [7].

Table 2 shows the performance comparison results between DeEPsnap and the other compared models across 10-fold cross-validation. From Table 2 we can see that the AUROC scores of all the compared models are above 91%, which indicates that the features extracted or learned from the five omics data are very effective for predicting human gene essentiality. DeEPsnap outperforms all the compared models, especially DNN, for all the five measures, which tells us that the proposed snapshot ensemble mechanism is a useful technique for improving model performance without incurring extra training cost. The AUPRC scores of three DNN-based models (DeEPsnap, DeepHE, DNN) are higher than those of the other compared models, which indicates that multilayer perceptron enhanced with deep learning techniques coupled with class weight can be more effective for coping with imbalanced learning problems. The comparison experiments also show that the training times of all the compared models but GAT are comparable (less than 16 minutes for 10-fold cross-validation on a laptop using CPU), while GAT needs more than 9 hours which is 35 times that of DeEPsnap. DeEPsnap only needs 50 epochs in training for each fold, but GAT needs about 2500 epochs (1500 to 3000 epochs across the 10-fold cross-validation).

**Table 2.** Performance comparison of DeEPsnap, DNN, DeepHE, SVM, RF, Adaboost, and GAT.

| Method | AUROC | AUPRC | BA | F1 | Accuracy |
| --- | --- | --- | --- | --- | --- |
| DeEPsnap | **0.9610 ± 0.0065** | **0.9382 ± 0.0113** | **0.8909 ± 0.0132** | **0.8062 ± 0.0196** | **0.9221 ± 0.0081** |
| DNN | 0.9511 ± 0.0062 | 0.9232 ± 0.0088 | 0.8730 ± 0.0120 | 0.7850 ± 0.0157 | 0.9152 ± 0.0062 |
| DeepHE | 0.9355 ± 0.0072 | 0.8953 ± 0.0148 | 0.8447 ± 0.0138 | 0.7467 ± 0.0246 | 0.9016 ± 0.0105 |
| SVM | 0.9562 ± 0.0097 | 0.8527 ± 0.0249 | 0.8883 ± 0.0205 | 0.7887 ± 0.0285 | 0.9123 ± 0.012 |
| RF | 0.9502 ± 0.0092 | 0.8532 ± 0.0206 | 0.7997 ± 0.0138 | 0.7268 ± 0.0239 | 0.9103 ± 0.0072 |
| Adaboost | 0.9460 ± 0.0078 | 0.8428 ± 0.0219 | 0.8417 ± 0.0116 | 0.7582 ± 0.0196 | 0.9102 ± 0.0076 |
| GAT | 0.9171 ± 0.0216 | 0.8554 ± 0.0360 | 0.8027 ± 0.0885 | 0.6568 ± 0.1255 | 0.8710 ± 0.0217 |

### Ablation study

In order to know the contribution of each type of feature used in DeEPsnap, we also evaluate it by removing one type of feature each time. The experiments use the same settings except for the input features. Table 3 gives the performance comparison results of DeEPsnap with different types of input features, which tells us that DeEPsnap with the integration of all the five types of features performs best which further confirms the contribution and complementarity of these five types of features. By removing one type of feature each time, we can evaluate the contribution of each type of feature to the overall performance and its complementarity to the other types of features.

**Table 3.**
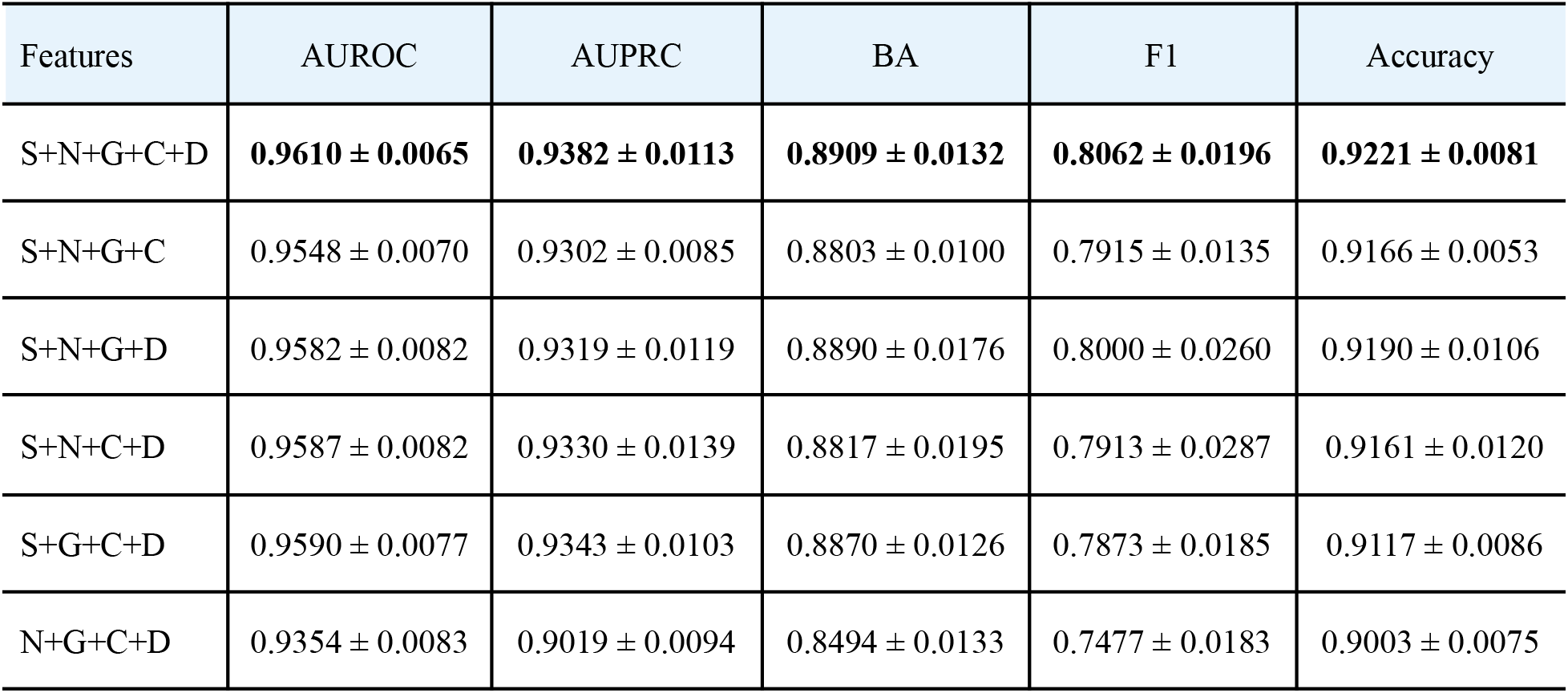
Performance comparison of DeEPsnap with different types of features. S: sequence features; N: network embedding features; G: gene ontology features; C: protein complex features; D: protein domain features.

| Features | AUROC | AUPRC | BA | F1 | Accuracy |
| --- | --- | --- | --- | --- | --- |
| S+N+G+C+D | **0.9610 ± 0.0065** | **0.9382 ± 0.0113** | **0.8909 ± 0.0132** | **0.8062 ± 0.0196** | **0.9221 ± 0.0081** |
| S+N+G+C | 0.9548 ± 0.0070 | 0.9302 ± 0.0085 | 0.8803 ± 0.0100 | 0.7915 ± 0.0135 | 0.9166 ± 0.0053 |
| S+N+G+D | 0.9582 ± 0.0082 | 0.9319 ± 0.0119 | 0.8890 ± 0.0176 | 0.8000 ± 0.0260 | 0.9190 ± 0.0106 |
| S+N+C+D | 0.9587 ± 0.0082 | 0.9330 ± 0.0139 | 0.8817 ± 0.0195 | 0.7913 ± 0.0287 | 0.9161 ± 0.0120 |
| S+G+C+D | 0.9590 ± 0.0077 | 0.9343 ± 0.0103 | 0.8870 ± 0.0126 | 0.7873 ± 0.0185 | 0.9117 ± 0.0086 |
| N+G+C+D | 0.9354 ± 0.0083 | 0.9019 ± 0.0094 | 0.8494 ± 0.0133 | 0.7477 ± 0.0183 | 0.9003 ± 0.0075 |

From Table 3, we can see that the combination of N + G + C + D performs worse which might be due to the fact that the three types of features in this combination utilize the PPI network topological information so that they have less complement effect. The other four combinations with sequence features perform only slightly worse than using all five types of features, which tells us that sequence features have a big complementarity with the other types of features. As shown in [7], when only using one type of feature, a deep model using network embedding features performs better than that using sequence features. Therefore, the above phenomenon doesn’t indicate that sequence features are superior to network features, but that sequence features are more complementary to the other three types of features.

### Enrichment analysis for essential genes

The list of essential genes was submitted to Enrichr [38] for analysis for GO Biological processes [34,35] and Reactome pathway [39] enrichment. The genes were highly statistically enriched for essential macromolecular biosynthetic processes as shown in Fig 4. These processes include translation, gene expression, ribosome biogenesis, mRNA processing, and more. The pathways enriched were similar to the GO term analysis with the exception that there is higher enrichment for cell cycle genes which is clearly an essential process. From this analysis, we conclude that our algorithm is selecting the types of genes and processes that would be expected for essential genes. This further confirms the fidelity of our method.

**Fig 4.**
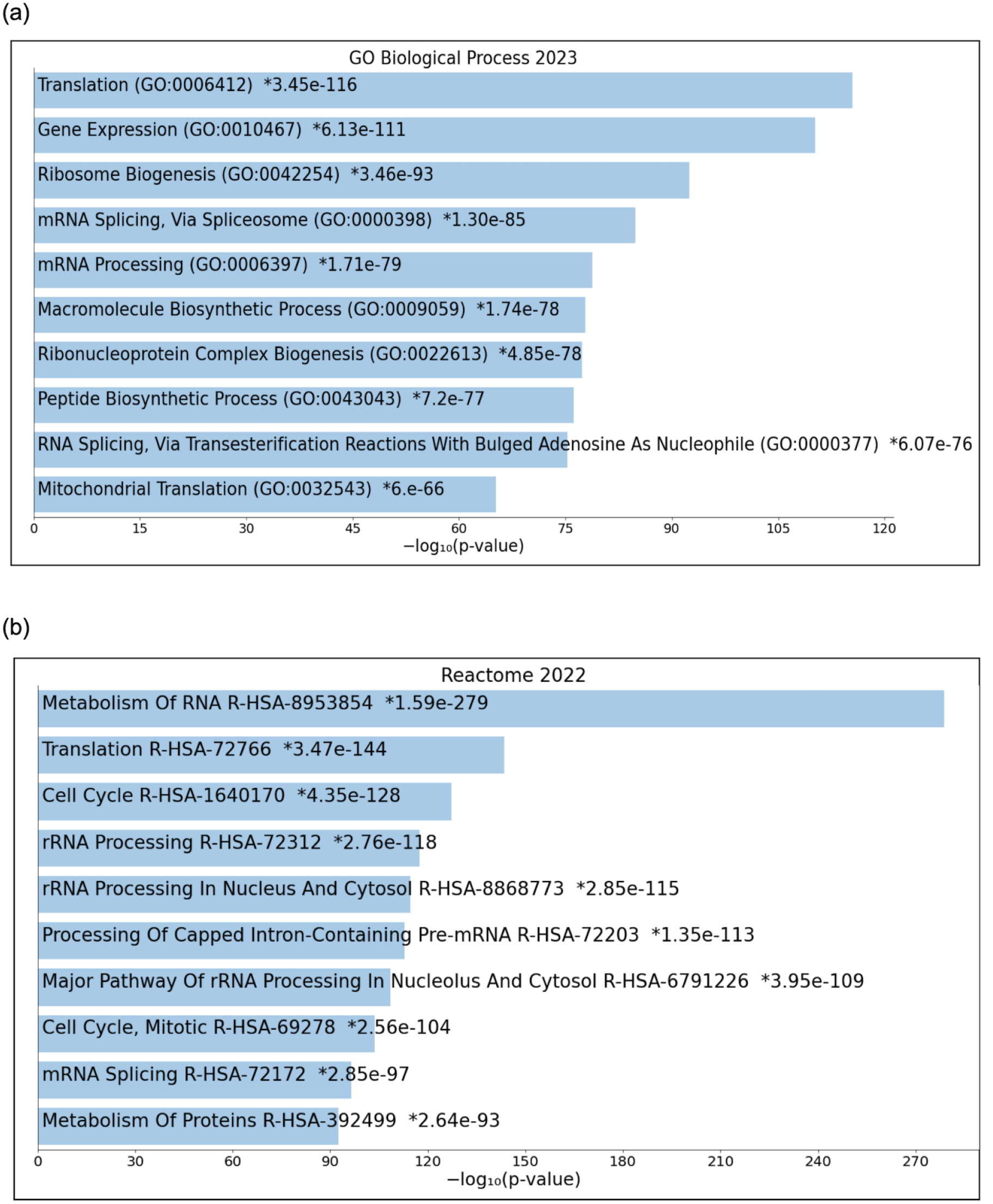
Biological process and pathway enrichment of the essential genes. (a) Bar plot of the top enriched BP terms; (b) Bar plot of the top enriched pathways in Reactome.

## Conclusions and future directions

In this paper, we propose a snapshot ensemble deep learning method, DeEPsnap, to predict human essential genes. DeEPsnap integrates five types of features extracted/learned from sequence and functional genomics data. It utilizes multiple deep-learning techniques and a cyclic annealing mechanism to train an ensemble of cost-sensitive classifiers to enhance the prediction accuracy of gene essentiality. Our 10-fold cross-validation experiments demonstrate: 1) the proposed snapshot ensemble deep learning method, DeEPsnap, is superior to the traditional machine learning models and some deep learning models, which is more effective for predicting human essential genes; 2) the extracted features from the five omics data are effective and complementary to gene essentiality prediction; 3) the proposed snapshot ensemble mechanism is promising for improving model’s prediction performance without incurring extra cost.

Although DeEPsnap is effective for predicting human essential genes, it still has some limitations. The most important one is that the current method cannot automatically extract useful features from four of the omics data used. Only features from the PPI network data are automatically learned based on a deep learning method, node2vec. The other four types of features are curated based on our understanding of the relationship between the data and gene essentiality. We believe that appropriate knowledge representations coupled with suitable feature learning algorithms can extract more useful features for predicting gene essentiality.

In the future, we are interested in how to use deep learning to automatically learn features from different types of biological data as well as to explore the appropriate knowledge representations for different omics data. For example, how to learn a low-dimension representation if we use all GO terms to encode genes instead of the subset of selected GO terms; it’s especially interesting to explore novel feature learning methods to extract more informative representation features from protein complex and protein domain data. In addition, exploring and integrating more biological data into the learning and classification model is another interesting direction, such as epigenomics data and gene expression profiles. Predicting cancer cell line-specific and tissue-specific essential genes by designing effective deep learning models and integrating cell line-specific and tissue-specific information would be very interesting, especially the prediction of essential genes across cell lines and tissues via useful transfer learning techniques. We are also interested in testing whether data editing [40] and clustering-aided techniques [41] are useful for imbalanced learning problems.

## Data availability

All data used in this study are third-party and freely accessible from public databases. Protein-protein interaction data are available from the BioGRID database at http://thebiogrid.org/download.php. Essential gene data and the corresponding sequence data from the DEG database are available at http://tubic.tju.edu.cn/deg/. DNA sequence and protein sequence data are available at https://useast.ensembl.org/Homo_sapiens/Info/Annotation. Gene Ontology data are available at http://current.geneontology.org/products/pages/downloads.html. Protein complex data are available at http://mips.helmholtz-muenchen.de/corum/. Protein domain data are from https://pfam.xfam.org/ and downloaded via Ensembl BioMart at http://useast.ensembl.org/info/data/biomart/index.html. The Python code of the model is freely available at https://github.com/wjxiao2020/DeEPsnap.

## References

[1] Wang T, Birsoy K, Hughes NW, Krupczak KM, Post Y, Wei JJ, et al. Identification and characterization of essential genes in the human genome. Science, 2015; 350(6264): 1096-101. doi: 10.1126/science.aac7041.

[2] Hart T, Chandrashekhar M, Aregger M, Steinhart Z, Brown KR, MacLeod G, et al. High-resolution CRISPR screens reveal fitness genes and genotype-specific cancer liabilities. Cell, 2015; 163(6): 1515–26. doi:10.1016/j.cell.2015.11.015.

[3] Grazziotin AL, Vidal NM, Venancio TM. Uncovering major genomic features of essential genes in Bacteria and a methanogenic Archaea. FEBS J. 2015; 282(17):3395–411. doi:10.1111/febs.13350.

[4] Liao BY, Zhang JZ. Mouse duplicate genes are as essential as singletons. Trends in Genetics. 2007; 23(8):378–81. doi:10.1016/j.tig.2007.05.006.

[5] Morgens DW, Deans RM, Li A, Bassik MC. Systematic comparison of CRISPR/Cas9 and RNAi screens for essential genes. Nature Biotechnology. 2016; 34(6):634–6. doi:10.1038/nbt.3567.

[6] Zhang X, Xu J, Xiao W-x. A New Method for the Discovery of Essential Proteins. PLoS ONE, 2013; 8(3): e58763. doi:10.1371/journal.pone.0058763.

[7] Zhang X, Xiao WX, Xiao WJ. DeepHE: Accurately predicting human essential genes based on deep learning. PLoS Comput Biol. 2020, 16(9): e1008229. doi:10.1371/journal.pcbi.1008229.

[8] Zhang X, Xiao W, Acencio ML, Lemke N, Wang X. An ensemble framework for identifying essential proteins. BMC Bioinformatics. 2016; 17:322. doi:10.1186/s12859-016-1166-7.

[9] Luo J, Qi Y. Identification of Essential Proteins Based on a New Combination of Local Interaction Density and Protein Complexes. PLoS ONE, 2015; 10(6): e0131418. doi:10.1371/journal.pone.0131418.

[10] Zhang X, Xiao W, Hu X. Predicting essential proteins by integrating orthology, gene expressions, and PPI networks. PLoS ONE, 2018; 13(4): e0195410. doi:10.1371/journal.pone.0195410.

[11] Li G, Li M, Wang J, Wu J, Wu F, Pan Y. Predicting essential proteins based on subcellular localization, orthology and PPI networks. BMC Bioinformatics. 2016; 17(Suppl 8):279. doi:10.1186/s12859-016-1115-5.

[12] Xiao W, Yan X, Zhang X. Pavement distress image automatic classification based on density-based neural network. International Conference on Rough Sets and Knowledge Technology (RSKT), July 2006; 686-692.

[13] Ozturk H, Ozgur A, Ozkirimli E. DeepDTA: deep drug-target binding affinity prediction. Bioinformatics, 34, 2018; i821–i829. doi: 10.1093/bioinformatics/bty593.

[14] Zhang X, Zhao D, Chen L, Min W. Batch mode active learning based multi-view text classification. The sixth international conference on fuzzy systems and knowledge discovery, August 2009; 7: 472–476.

[15] Xiao W, Zhang X. Active transductive KNN for sparsely labeled text classification. The 6th international conference on soft computing and intelligent systems, jointly the 13^th^ international symposium on advanced intelligence systems. November 2012; 2178-2182.

[16] Guo F, Dong C, Hua H, Liu S, Luo H, Zhang H, et al. Accurate prediction of human essential genes using only nucleotide composition and association information. Bioinformatics, 2017; 33(12), 1758–1764. doi: 10.1093/bioinformatics/btx055.

[17] Schapke J, Tavares A, Recamonde-Mendoza M. EPGAT: Gene Essentiality Prediction With Graph Attention Networks. IEEE/ACM Transactions on Computational Biology and Bioinformatics, 2022; 19(3): 1615–1626. doi: 10.1109/tcbb.2021.3054738.

[18] Wang J, Zhang L, Sun J, Yang X, Wu W, Chen W, et al. Predicting drug-induced liver injury using graph attention mechanism and molecular fingerprints. Methods, 2024; 221: 18–26. doi: 10.1016/j.ymeth.2023.11.014.

[19] Wang T, Sun J, Zhao Q. Investigating cardiotoxicity related with hERG channel blockers using molecular fingerprints and graph attention mechanism. Computers in Biology and Medicine, 2023; 153(2023): 106464. doi: 10.1016/j.compbiomed.2022.106464.

[20] Gao H, Sun J, Wang Y, Lu Y, Liu L, Zhao Q, et al. Predicting metabolite-disease associations based on auto-encoder and non-negative matrix factorization. Briefings in Bioinformatics, 2023; 24(5): 1–13. doi:10.1093/bib/bbad259.

[21] Zhang X, Acencio ML and Lemke N. Predicting Essential Genes and Proteins Based on Machine Learning and Network Topological Features: A Comprehensive Review. Front. Physiol. 2016; 7:75. doi:10.3389/fphys.2016.00075.

[22] Grover A, Leskovec J. node2vec: Scalable Feature learning from networks. KDD’16: Proceedings of the 22nd ACM SIGKDD International Conference on Knowledge Discovery and Data Mining. August 2016;855-864. doi:10.1145/2939672.2939754.

[23] Zeng M, Li M, Wu FX, Li YH, Pan Y. DeepEP: a deep learning framework for identifying essential proteins. BMC Bioinformatics 2019; 20(Suppl 16):506. doi: 10.1186/s12859-019-3076-y.

[24] Zeng M, Li M, Fei Z, Wu F, Li Y, Pan Y, et al. A deep learning framework for identifying essential proteins by integrating multiple types of biological information. IEEE/ACM Transactions on Computational Biology and Bioinformatics. 2019 Feb 5. doi: 10.1109/TCBB.2019.2897679.

[25] Zhang X, Xiao W, Xiao W. A deep learning framework for predicting human essential genes by integrating sequence and functional data. bioRxiv, 2020. doi: 10.1101/2020.08.04.236646.

[26] Hasan MA, Lonardi S. DEEPLYESSENTIAL: A deep neural network for predicting essential genes in microbes. BioRxiv, 2019. doi: 10.1101/607085.

[27] Li Y, Zeng M, Zhang F, Wu FX, Li M. DeepCellEss: cell line-specific essential protein prediction with attention-based interpretable deep learning. Bioinformatics, 2023; 39(1): btac779. doi: 10.1093/bioinformatics/btac779.

[28] Yue Y, Ye C, Peng PY, Zhai HX, Ahmad I, Xia C, et al. A deep learning framework for identifying essential proteins based on multiple biological information. BMC Bioinformatics. 2022; 23:318. doi: 10.1186/s12859-022-04868-8.

[29] Blomen VA, Májek P, Jae LT, Bigenzahn JW, Nieuwenhuis J, Staring J, et al. Gene essentiality and synthetic lethality in haploid human cells. Science. 2015; 350(6264):1092-6. doi: 10.1126/science.aac7557.

[30] Huang G, Li Y, Pleiss G, Liu Z, Hopcroft JE, Weinberger KQ. Snapshot Ensembles: Train1, get M for free. 5th International Conference on Learning Representations, April 24-26, 2017; ICLR 2017, France.

[31] Luo H, Lin Y, Gao F, Zhang CT, Zhang R. DEG 10, an update of the database of essential genes that includes both protein-coding genes and noncoding genomic elements. Nucleic Acids Res. Jan 2014; 42 (Database issue): D574–80. doi: 10.1093/nar/gkt1131.

[32] Ruffier M, Kähäri A, Komorowska M, Keenan S, Laird M, Longden I, et al. Ensembl core software resources: storage and programmatic access for DNA sequence and genome annotation. Database. 2017. doi: 10.1093/database/bax020.

[33] Stark C, Breitkreutz BJ, Reguly T, Boucher L, Breitkreutz A, Tyers M. Biogrid: A General Repository for Interaction Datasets. Nucleic Acids Res. Jan 1, 2006; 34: D535–9.

[34] Ashburner, et al. Gene ontology: tool for the unification of biology. Nat Genet. May 2000;25(1):25–9.

[35] The Gene Ontology Consortium. The Gene Ontology knowledgebase in 2023. Genetics. 2023 May 4; 224(1): iyad031. doi: 10.1093/genetics/iyad031

[36] Giurgiu M, Reinhard J, Brauner B, Dunger-Kaltenbach I, Fobo G, Frishman G, et al. CORUM: the comprehensive resource of mammalian protein complexes-2019. Nucleic Acids Res. 2018 Oct 24. doi: 10.1093/nar/gky973.

[37] El-Gebali S, Mistry J, Bateman A, Eddy SR, Luciani A, Potter SC, et al. The Pfam protein families database in 2019. Nucleic Acids Research. January 2019; 47(D1): D427–D432. doi:10.1093/nar/gky995.

[38] Kuleshov MV, Jones MR, Rouillard AD, Fernandez NF, Duan Q, Wang Z, et al. Enrichr: a comprehensive gene set enrichment analysis web server 2016 update. Nucleic Acids Research. 2016; gkw377.

[39] Milacic M, Beavers D, Conley P, Gong C, Gillespie M, Griss J, et al. The Reactome Pathway Knowledgebase 2024. Nucleic Acids Research. 2024. doi: 10.1093/nar/gkad1025.

[40] Zhang X, Xiao W. Active semi-supervised framework with data editing. Computer Science and Information Systems. 2012; 9(4): 1513–1532. doi: 10.2298/CSIS120202045Z.

[41] Zhang X, Xiao W. Clustering based two-stage text classification requiring minimal training data. Computer Science and Information Systems. 2012; 9(4):1627–1643. doi: 10.2298/CSIS120130044Z.

